## Supplementary Information for "Diffusion controls local versus dispersed inheritance of histones during replication and shapes epigenomic architecture"

**Contents**

| Section | Page |
|---|---|
| 1. Detailed description of Model 1 and assumptions | 1 |
| 2. Detailed description of Model 2 and assumptions | 3 |
| 3. Description of data pre-processing | 5 |
| 4. The bioinformatics pipeline | 6 |
| 5. Supplementary Figures and Legends | 7 |
| 6. Supplementary references | 12 |

### 1. Detailed description of Model 1 and assumptions.

*Defining the model:* To model the process of DNA replication, we first define a two-dimensional "parent" matrix $\underline{P}$, representing the parent cell undergoing replication. The rows of $\underline{P}$, indexed by $i$ ($i \in 1, 2, \ldots, R$), represent $R$ different regions or loci within the same or different chromosomes. The columns of $\underline{P}$, indexed by $j$ ($j \in 1, 2, \ldots, N$), represent $N$ nucleosomal sites along each genomic loci. $\underline{P}(i, j)$ is initialized with the value "1" denoting the presence of a histone at each site $j$ within the locus $i$. Only along one special row denoted by $i = R^*$ (representing for example the *Hoxc6* locus), are the "1"s followed by a "*". "1*" represents parental histones tagged with biotin, in accordance with the experimental design of Escobar et al.

Next, two daughter matrices $\underline{D_1}$ and $\underline{D_2}$ of dimension $R$ x $N$ each are initialized with "0"s as entries; the rows of these matrices describe the newly formed naked DNA strands (without any histones) for each loci. Note that $\underline{D_1}$ and $\underline{D_2}$ represent the two daughter cells of the parent $\underline{P}$, so row $i$ of each daughter represents the same chromosomal locus $i$ in the parent $\underline{P}$. Finally, physical distances between nucleosomal sites in one column of $\underline{P}$ and sites in the same column of $\underline{D_1}$ or $\underline{D_2}$ are defined based on a simple rule: $x_{i,i'} = |i - i'|$, for any $j$, where $i$ represents the row number of $\underline{P}$, $i'$ the row number of $\underline{D_1}$ (or $\underline{D_2}$) and $j$ denotes the column number of $\underline{P}$.

The processes of replication fork progression and consequent histone transfer between $\underline{P}$ and $\underline{D_1}$ or $\underline{D_2}$ are modeled by first choosing column $j = 1$ of $\underline{P}$ and probabilistically moving all its entries ("1" or "1*") to sites in the first column of either $\underline{D_1}$ or $\underline{D_2}$. This process is then repeated serially for all the columns $j = 2, \ldots, N$. The movement of entries in any column of $\underline{P}$ to a randomly chosen row in the same column of either $\underline{D_1}$ or $\underline{D_2}$ is carried out via a probabilistic rule to model diffusion among the positions of this column:

$$p_{i,i'} = \frac{1}{\sqrt{4\pi DT}} e^{-\frac{x_{i,i'}^2}{4D}T}, \qquad (1)$$

where $p_{i,i'}$ is the probability of a histone in row $i$ of $\underline{P}$ moving to row $i'$ of $\underline{D_1}$ or $\underline{D_2}$ (within the same column $j$), $D$ is the diffusion constant and $T$ represents some characteristic time over which the diffusion can happen. For this entire work, $T$ was set to 1 for simplicity. For simulations of loci in active genomic regions, the diffusion constant $D$ is larger than that in repressed regions, and is obtained from fits to experimental data from Escobar et al.

Since the redistribution rule is based on diffusion given by Eq. (1), sites on the daughters that are closer to the parental histone site (smaller values of $x_{i,i'}$) are more likely to be destination spots for the dislodged histone; sites with larger $x_{i,i'}$ values have small but non-zero probability of receiving the histone. Once all entries of all columns of $\underline{P}$ have been transferred to $\underline{D_1}$ and $\underline{D_2}$, sites in the two daughters that remain empty are filled up with new "1"s, representing newly synthesized histones. Note that this process keeps track of where the "1*" tagged histones land in the daughters. This series of steps represents the completion of one cell cycle event. We then randomly (with probability 0.5) choose one of the two daughter matrices $\underline{D_1}$ or $\underline{D_2}$ and assign it to be the parent $\underline{P}$, and the whole procedure is repeated many times to mimic multiple cell division events. Over successive cell divisions, we keep track of the fraction of columns corresponding to the row $R^*$ which retain the "1*" entry, mimicking tracking of tagged histone dilution at specific gene loci in the experiments of Escobar et al. Due to the possibility of 1* histones diffusing to rows other than $R^*$ in the daughters, the number of tagged histones in this particular row decreases over time.

*Assumptions underlying Model 1:* There are naturally a number of assumptions that were implicitly or explicitly made in defining Model 1 above. Here we state and discuss each of these assumptions.

(a) We use a 1D diffusion equation for simplicity and the lattice model has distances only in 1D. This is an oversimplification of the full 3D problem. However, in higher dimensions the effects of diffusion would be more rather than less, since more sites would be available for parental histones to diffuse to. The more limiting result of this simplification is that currently it is not possible for us to quantitatively compare diffusion constants we extract from fits to experiments with real diffusion constants reported in the literature.

(b) Another assumption implicit in Model 1 is that only the histones diffuse. In reality, DNA fluctuations are also important[1] and hence an "effective" diffusion constant combining histone as well as DNA diffusion is likely to be more relevant. However, this assumption is not too limiting since $D$ in our model could be interpreted as a combination of both.

(c) Redistributing histones one column at a time ("vertical" diffusion within a column is allowed, but not "horizontal" diffusion along rows) translates to a simplifying assumption that a dislodged set of histones must find their new spots before the next set of parental histones get dislodged. This may be a reasonable assumption, since a number of studies in the past have demonstrated that nucleosome positioning is rapidly re-established behind the replicating fork[2] . Note that this assumption does not preclude diffusion from one locus to another locus on the same chromosome – as mentioned previously in the model definition, different rows in the $\underline{P}$, $\underline{D_1}$ and $\underline{D_2}$ matrices can be interpreted as loci either on the same or on different chromosomes. However, it would be interesting to investigate potential scenarios where replication fork progression occurs faster than the time in which dislodged histones can reassemble behind the fork. In such situations, "horizontal" diffusion will also contribute to the redistribution of histones across loci. Such studies are left for future work.

(d) Whether free histones (i.e. histones not associated with DNA in the nucleosomal complex) exhibit unhindered diffusion or sub-diffusion is not clear. Salt induced disassembled nucleosomes have been demonstrated to exhibit free diffusion[3], hence for simplicity we have used a model of free diffusion in this work. While sub-diffusive models of histone dynamics may change quantitative aspects of our results, qualitatively we would not expect differences in the predicted histone mark similarity patterns.

(e) Another limitation of our model and results is that since Hi-C provides correlates of physical proximity and not real distances, it is challenging to map our distance $x$ in simulations to the spatial proximity measure in Hi-C datasets. This limitation also currently prevents us from estimating how large the DADs could be and how the DAD sizes differ between active and repressed chromatin regions. Methods to infer physical distances from Hi-C contact maps are being currently developed, and might help resolve these intriguing questions in future studies. Indeed, since the slope of the median $Q(i,j)$ vs $x$ plot is directly related to the diffusion constant, estimating the diffusion constants in various genomic regions from these plots presents an exciting possibility for the future.

**2. Detailed description of Model 2 and assumptions.**

*Defining the model:* Model 1 was designed to study the kinetics of histone dilution due to cell division and diffusion. Model 2 is a modified version of the previous model, to allow investigation of histone modification patterns resulting from diffusion and replication. To include histone modifications, we re-interpret the "1"s in the parent matrix $\underline{P}$ of Model 1 to be histones with a particular modification of interest (say H3K27ac), and "0"s to be newly synthesized histones without any parental marks. We include an additional label "2" to denote histones with marks other than the one of interest. This is done to maintain similarity with our bioinformatics procedure (to be discussed later; see Section **S4**), where we look at histone modification signals in coarse grained chromosome loci where a variety of histone marks are present. We also modify the description of $\underline{P}$ by assigning two labels to the rows: the first $R/2$ rows are labelled "$E$", denoting loci from early replicating regions, while the last $R/2$ rows are labelled "$L$" to denote late replicating loci. Throughout the paper, we used $R = 20$ and $N = 200$. The precise values of $R$ and $N$ do not qualitatively change any of the results in this study.

To assign initial values (1's or 2's) to every element $(i, j)$ of $\underline{P}$, we first chose a random number from an exponential distribution with parameter 130, which represents the number of 1's in the first row. These 1's were then randomly assigned to the 200 columns of the first row, and the remaining columns were assigned the value 2. Thus every histone is assumed to have some modification – either 1 or 2. This procedure is then repeated for all the rows, thus assigning a modification to every element of the parent matrix $\underline{P}$. Once again, the choice of the exponential distribution parameter does not matter, and we checked that assigning different parameter values to the first $R/2$ rows corresponding to $E$ loci vs the last $R/2$ rows corresponding to $L$ loci also does not matter. Neither does the functional form of the distribution matter. The two daughter matrices $\underline{D_1}$ and $\underline{D_2}$ were initialized with 0's for every element, indicating DNA with no histones present.

As in Model 1, we assign distances to every pair of rows $(i, j)$, but with the following modification: row-pairs $(i, j)_{EE}$ where both the rows belong to $E$, have a larger average distance than $(i, j)_{LL}$ pairs where both rows belong to $L$. This is achieved by simply setting $x_{i,i'} = |i - i'| + 1$ whenever $(i, i')$ are $E$ loci. This ensures that the model is consistent with the fact that late replicating loci tend to lie in heterochromatin which is more compact than euchromatin, and hence are expected to have smaller inter-loci distances on average. The probabilities of histone mark re-distribution during cell division are assigned using Eq. (1) as before. The only difference now is that two distinct diffusion constants are used for $E - E$ and $L - L$ transitions: $D_{EE}$ and $D_{LL}$, where $D_{EE} > D_{LL}$. Results from Model 2 were generated with $D_{EE} = 1.2$ and $D_{LL} = 0.07$ throughout the main text. Qualitative results of this work do not depend on the precise values of these diffusion constants, as long as the inequality $D_{EE} > D_{LL}$ is maintained.

This modified model is then simulated for many cell cycles as described for Model 1, using Eq. (1) to probabilistically move both the histone marks ("1" and "2") to either one of the daughter matrices. Transitions within $E - E$ or $L - L$ rows differ simply in the use of different values of the diffusion constant. After a column of $\underline{P}$ gets redistributed between the columns of $\underline{D_1}$ and $\underline{D_2}$, a certain number of original 0's in the daughters will remain. This corresponds to histone mark dilution due to cell division. To prevent the histone mark levels from diminishing to zero due to this dilution phenomenon, we implement a local mark-copying rule to update the 0's to either 1 or 2 at the end of each cell cycle. We identify where all 0's in the daughters are located after a cell cycle is complete, and one of the two nearest neighbor's marks is randomly assigned to the empty site. If a 0 is flanked on both sides by 0's, then we look for the nearest non-0 mark, and convert all the 0's to that non-0 mark. If 0's occur at the edges of the matrices ($j = 1, N$), then the mark on $j = 2$ or $j = N - 1$ is copied over. This process is carried out till no more 0 sites remain in the daughters. Finally, a daughter is randomly selected between $\underline{D_1}$ and $\underline{D_2}$ and assigned to $\underline{P}$. $\underline{D_1}$ and $\underline{D_2}$ are re-initialized with 0's and the full cell cycle is repeated.

*Assumptions underlying Model 2:* Since Model 2 is an extension of Model 1, all the assumptions underlying the latter also hold true here. The major additional assumption in Model 2 is in the manner in which the mark copying is carried out. There is evidence to suggest that somewhat long-range cooperativity might be important in the mark-copying process[4,5], though the precise nature of this range and mechanisms of cooperativity are still not well understood. For simplicity, we only model nearest-neighbour interactions, where the marks on the immediate two neighbours of a site with a 0 mark affect the final state of the unmodified histone. However, including more complex mechanisms of mark copying should not qualitatively affect any of the results presented here.

**3. Description of data preprocessing.**

Data processing for sequencing datasets:

Replication sequencing dataset-

We sourced the replication timing dataset of the cell lines for early (S1) and late (S4) phase from ENCODE[6] (https://www.encodeproject.org/). The .bam files were converted to .bed files using the Bedtools bamtobed program. The .bed files were then intersected with a custom made .bed file (with HiC information- chromosome number, fragment start, fragment end, HiC bin number) to create 1-Mb fragments (HiC data resolution) of the replication sequencing data using Bedtools intersect program.

Defining early and late fragments: For 1 Mb segments that had exclusively late or early signal, the segments were directly classified as E or L respectively. However, many 1 Mb segments had both early and late signals. For such segments, we used the normalized signal values from the .bed files to classify coarse-grained Early and Late fragments in the following manner – for any 1 Mb fragment $i$, we first calculated the difference between the signal values (total early signal – total late signal). If the difference was positive (> cutoff), the fragment $i$ was assigned Early while if the difference was negative (< -cutoff), the fragment $i$ was assigned Late. The cutoff was used to classify the fragments with more certainty of being early or late; the central results of the paper do not depend on the precise value of the cutoff.

Note: For the lymphoblastoid cell line GM06990, we obtained the replication sequencing data from the lymphoblastoid cell line GM12878.

ChIP Sequencing dataset-

We sourced the ChIP sequencing dataset of the cell lines for different histone modifications from ENCODE. The .bed files were intersected with the custom made .bed file (with HiC information) to create 1-Mb fragments of the ChIP sequencing data using the bedtools program. This operation was performed in the R-script using the library bedr (uses Bedtools as dependency).
Note: For the lymphoblastoid cell line GM06990, we obtained the ChIP sequencing data from the lymphoblastoid cell line GM12878.

<u>HiC Sequencing dataset-</u>
We sourced the HiC dataset (correlation matrices) of the cell lines from Gene Expression Omnibus (GSE18199)[7]. The data was already pre-processed and ready to be integrated into our analysis. Details of the processing of the HiC data can be found in the methods & the supplementary section of Lieberman et al[7].

**4. The bioinformatics pipeline.**
For a given chromosomal comparison between chromosome A and chromosome B, where A, B $\in \{1, 2, \dots, 22\}$, we carry out the following steps:

1. We read in the .bed file from the ChIP sequencing data (see Section 3 for details) and subset this file to keep only chromosome A and B.
2. We read in the .bed files (early and late) from the replication sequencing data (see Section 3 for details) and subset these files to keep only chromosome A and B.
3. Next, we read in the HiC data (HiC matrix) for the 2 given chromosomes. In the matrix, each correlation value corresponds to a comparison between fragment $i$ and $j$ of chromosome A and B. We subset the matrix and keep only those $(i, j)$ pairs where $i$ and $j$ belong either to early or late classifications. We stored these correlation values for plotting the results.
4. Finally, we calculated the $Q(i, j)$ value using the formula for a given pair of early/ late fragments (see Methods section). We repeated this procedure for all such early-early and late-late pairs and until all chromosomal pairs have been analyzed (intrachromosomal & interchromosomal). We stored all $Q(i, j)$ values to plot the results.

### 5. Supplementary Figures and Legends

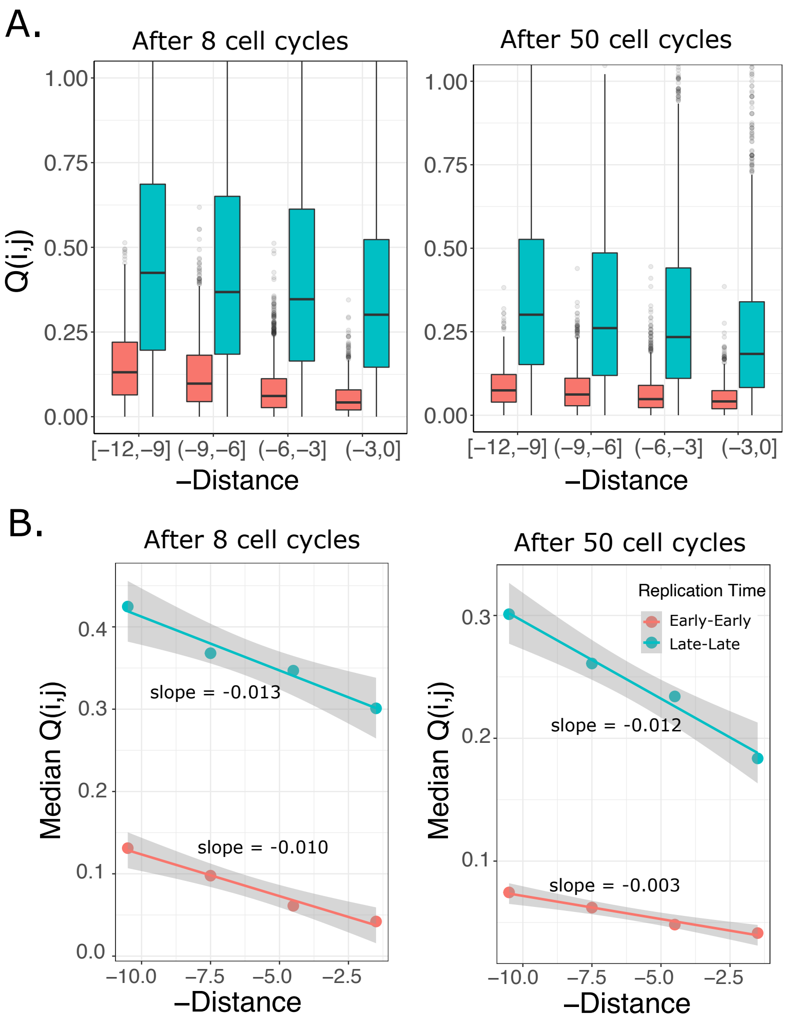


**Supplementary Figure 1:** Simulations with different starting histone modification signals between E and L loci do not qualitatively affect the central results. Every row of the parent matrix was initialized with a draw from an exponential distribution with parameter 130, for E loci, and parameter 20 for L loci. (A) The $Q(i,j)$ patterns after 8 and 50 cell cycles and (B) the slopes of the distance dependence of $Q(i,j)$ show the same expected patterns as shown in Fig. 3 of the Main Text.

Fig. S2

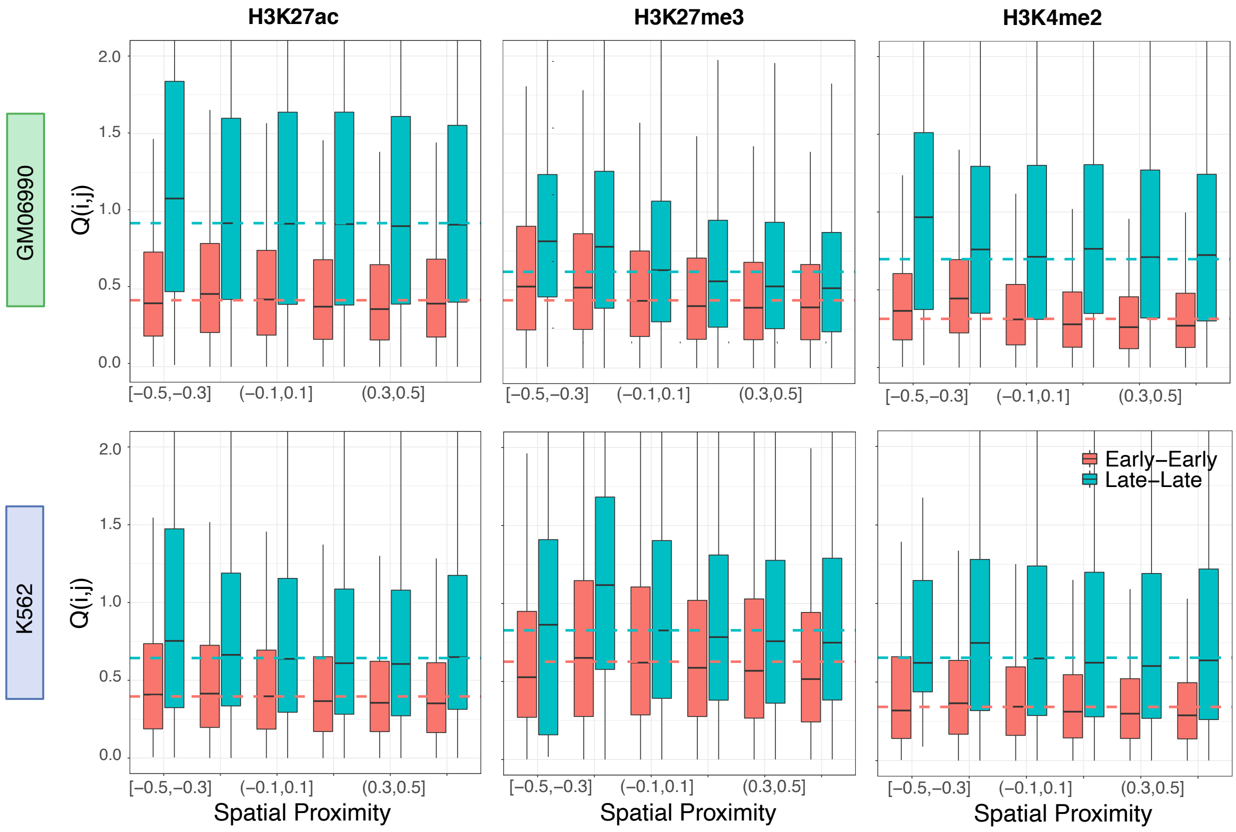


**Supplementary Figure 2:** Histone mark similarity patterns are unchanged for a cutoff value of 500 for the Early vs Late assignment. Top row shows the $Q(i,j)$ patterns for three marks H3K27ac, H3K27me3 and H3K4me2 for the GM06990 cell line, while the bottom row is the same for the K562 cell line.

Fig. S3

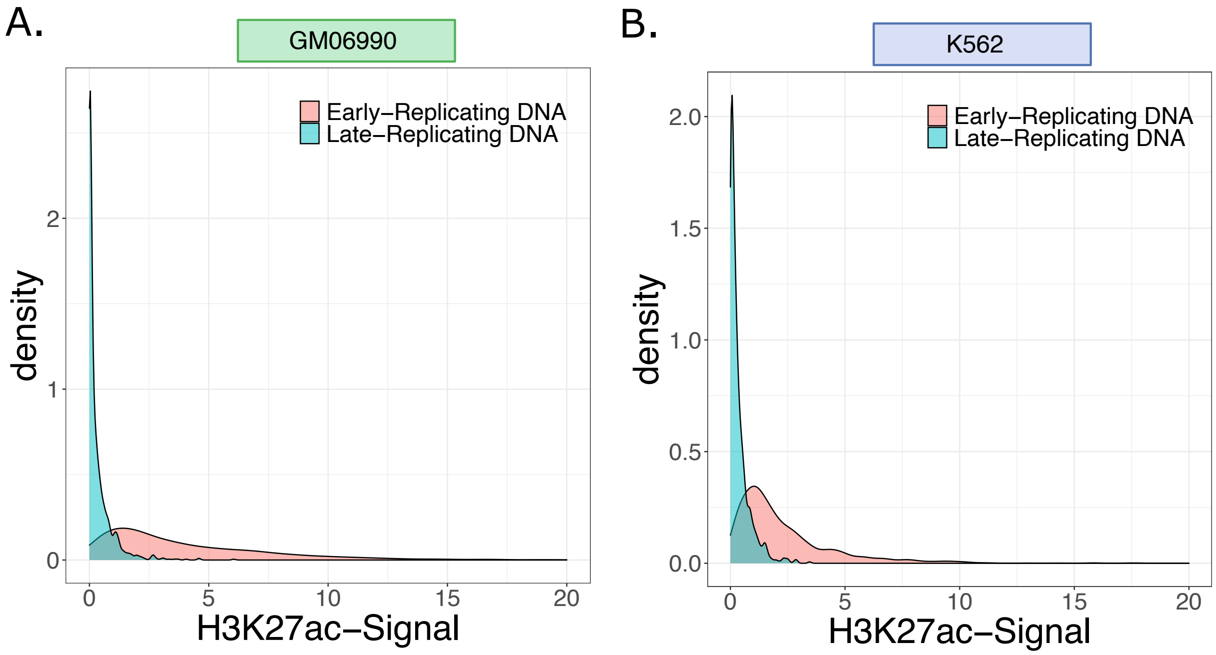


**Supplementary Figure 3:** An example showing the difference in histone modification levels between early and late loci, for the H3K27ac mark. Cutoff for Early-Late classification here was 1000. (A) GM06990 cell line and (B) K562 cell line.

Fig. S4

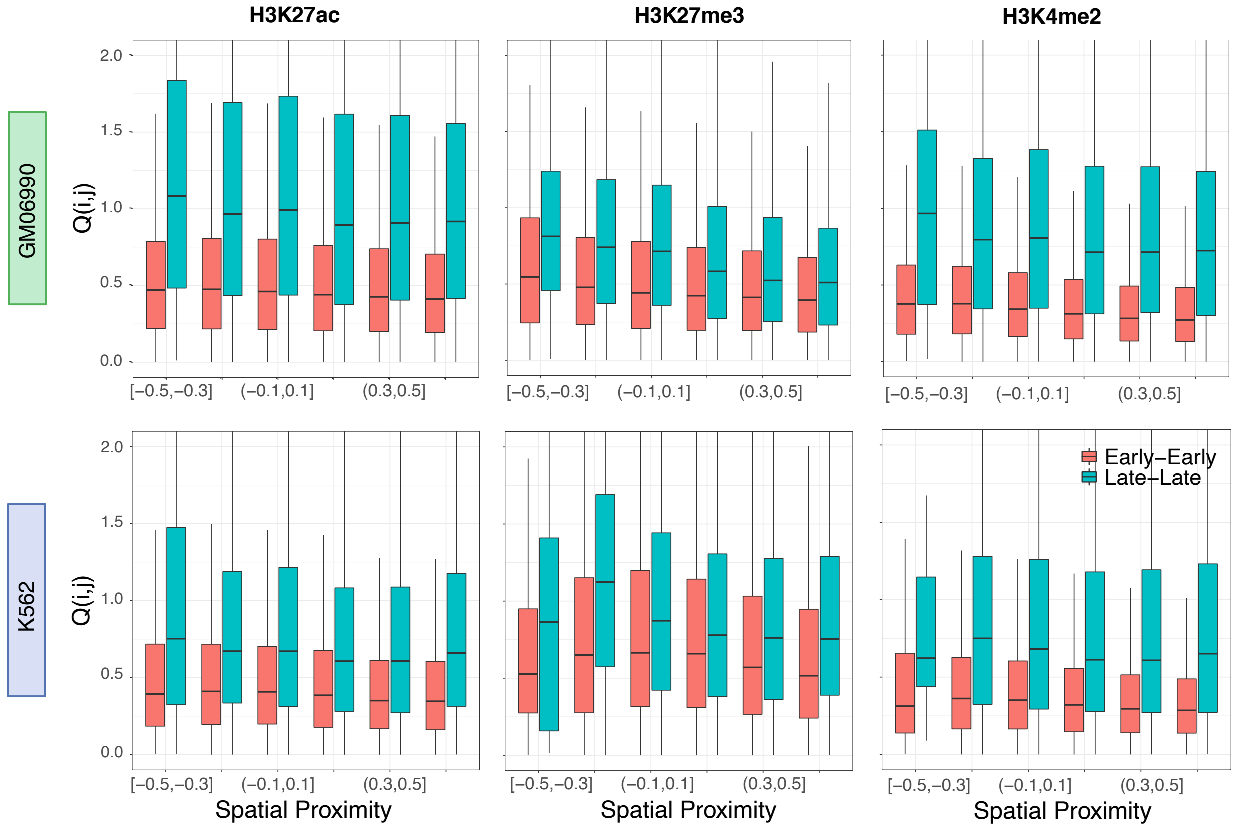


**Supplementary Figure 4:** *Intra*-chromosomal $Q(i,j)$ patterns show all the diffusion-induced patterns. Top row shows the $Q(i,j)$ patterns for three marks H3K27ac, H3K27me3 and H3K4me2 for the GM06990 cell line, while the bottom row is the same for the K562 cell line.

Fig. S5

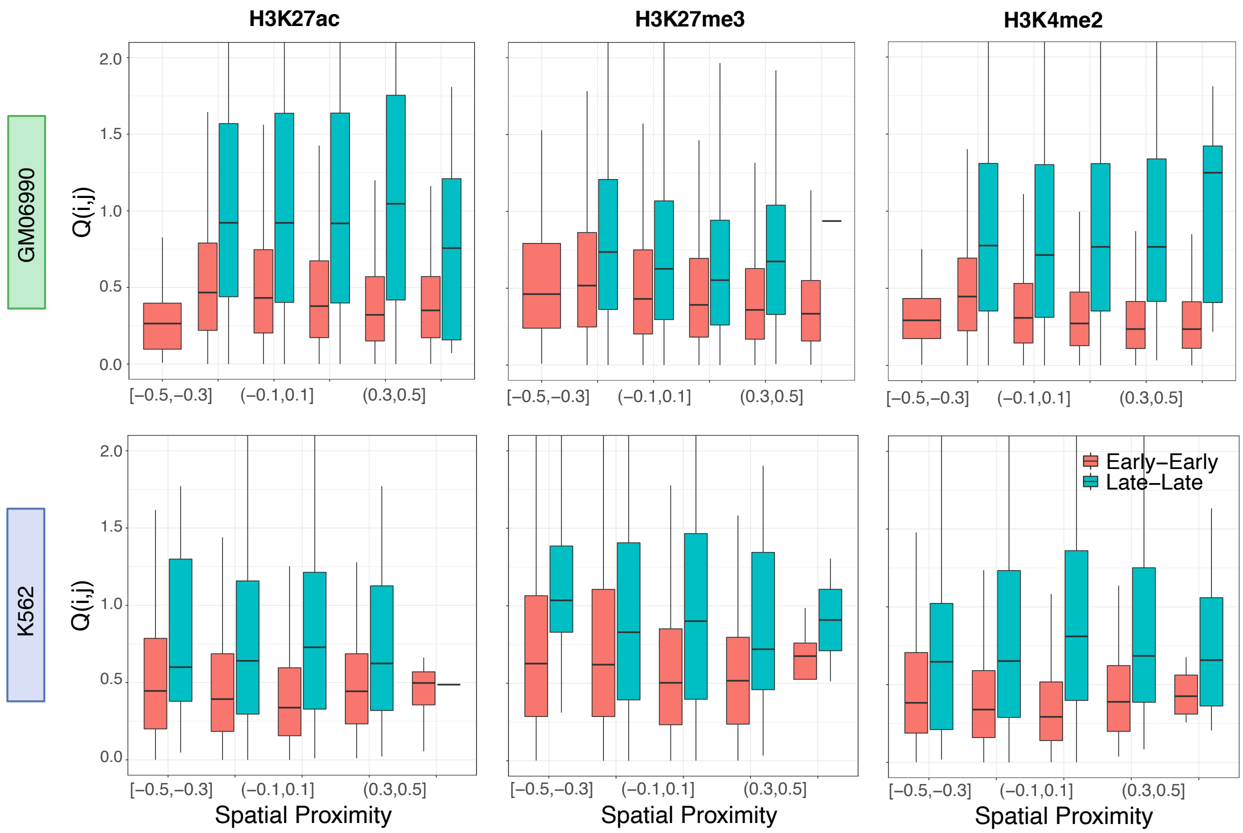


**Supplementary Figure 5:** *Inter*-chromosomal $Q(i,j)$ patterns show the diffusion-induced patterns for some marks, mainly in the GM06990 cell line. Top row shows the $Q(i,j)$ patterns for three marks H3K27ac, H3K27me3 and H3K4me2 for the GM06990 cell line, while the bottom row is the same for the K562 cell line. The distance dependence is clearer for the Early-Early loci, consistent with the expectation that diffusion is easier in active genomic loci as compared to repressed loci.

Fig. S6

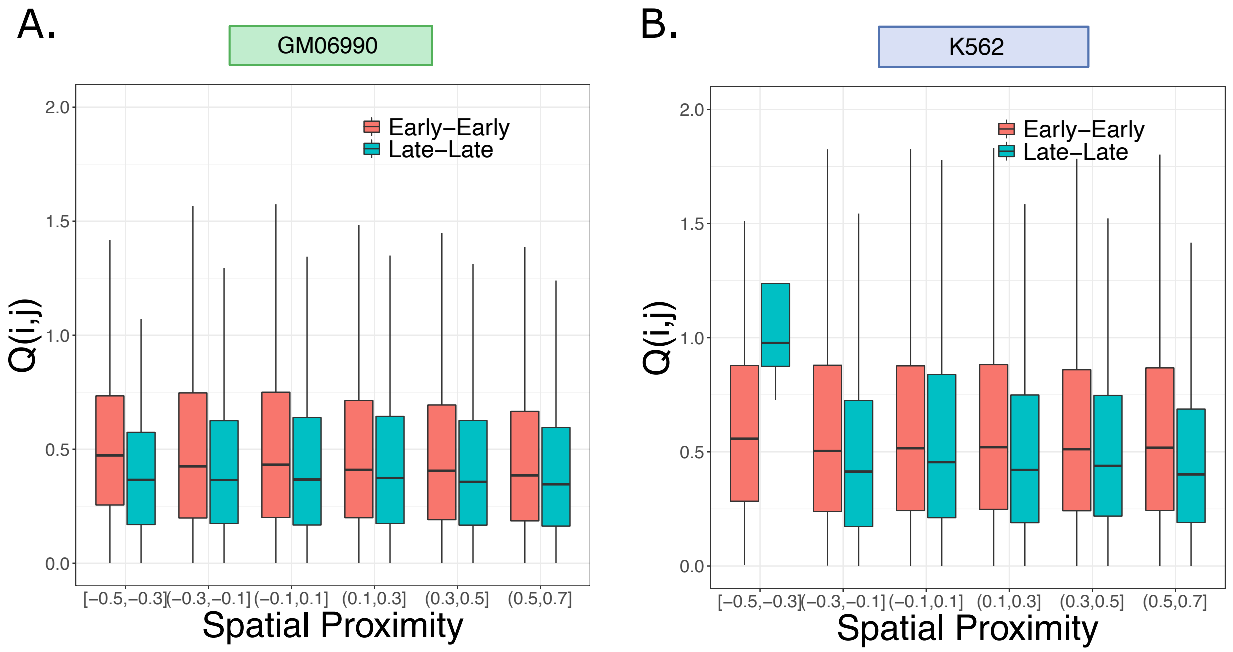


**Supplementary Figure 6:** *Intra*-chromosomal $Q(i,j)$ patterns for the H3K9me3 histone mark, in (A) the GM06990 and (B) the K562 cell lines. None of the diffusion-induced patterns can be observed for this mark, consistent with the idea that little to no diffusion mediated redistribution occurs within constitutive heterochromatin regions, even along the same chromosome.
